# Pepid: a Highly Modifiable, Bioinformatics-Oriented Peptide Search Engine

**DOI:** 10.1101/2023.10.30.564469

**Authors:** Jeremie Zumer, Sébastien Lemieux

**Affiliations:** Institut de Recherche en Immunologie et Cancerologie, Quebec, Canada; Department of Biochemistry and Molecular Medicine, Université de Montréal, Quebec, Canada

**Keywords:** Deep Learning, Proteomics, Peptide Identification, Open Source

## Abstract

**Motivation:** Current peptide search engines are optimized for wet-lab workflows, i.e. they operate in an “end-to-end” manner to achieve good identification results, not to be modified or provide algorithmic insight. This makes developing new software methods to solve problems in peptide identification methods difficult, often requiring a full engine rewrite. Recently, many deep learning methods were proposed as solutions to various parts of the peptide identification task, but virtually none of those methods have been implemented in any actual peptide search process. We believe that the lack of a reliable bioinformatics research platform for peptide identification that enables such integrations is slowing down proteomics research as a whole.

**Results:** We present pepid, a bioinformatics research-oriented peptide search engine. Unlike other search engines, pepid is specifically designed with ease of computational research in mind. Our design is highly flexible and allows easy modifications with little required software development expertise, allowing researchers to focus on analysing and improving peptide identification methods.It also takes recent computational trends into account, such as the recent slew of deep learning publications in proteomics, and features a multi-phased batched operations design that is more appropriate than the spectrum batch “end-to-end” designs of existing search engines for those approaches. We show that pepid is competitive with common engines in terms of both identification rates and runtime, forming a minimum required baseline to enable further identification research.

**Availability and Implementation:** Pepid is available as open source software under the MIT license at https://github.com/lemieux-lab/pepid. Other data referenced in the text is 3rd party. The selected yeast proteome can be found on SwissProt with accession ID UP000002311 while the human proteome’s accession ID is UP0000005640. The ProteomeTools spectra can be found in the PRIDE archive under accession ID PXD004732 and the One Hour Yeast Proteome can be found at the ChorusProject at https://chorusproject.org/anonymous/download/experiment/-8823069691100997209 and https://chorusproject.org/anonymous/download/experiment/449795368199176159.

## 1. Introduction

Peptide identification is a technically and scientifically difficult problem both on the wet- and dry-lab side. Current peptide search engines are designed for “end-to-end” operations, i.e. to provide a streamlined process for proteomics experiments considering wet-lab users, with non-separable operations from spectrum and protein processing all the way to scoring and output. This has the unintended consequence of making computational research in peptide identification harder, because deviating from the established paradigms effectively requires rewriting the entire engine. Recently, many deep-learning algorithms have been proposed to solve various problems relevant to peptide identification, such as theoretical spectrum generation Gessulat et al., 2019, Liu et al., 2020], retention time prediction Zeng et al., 2022, Giese et al., 2021], peptide-spectrum scoring, and so on (we direct interested readers to more comprehensive reviews of deep learning methods in proteomics, such as Wen et al. 2020] and Meyer 2021]). Unfortunately, these methods cannot be readily tested on top of most labs’ software workflow because making the requisite changes requires good software development expertise, as well as time and financial budget that may not always seem like a good use of resources to most labs. As a result, it is not clear which of these methods, if any, is actually useful for database-driven peptide searches, and even in the rare cases where those methods are implemented with an existing search engine Wilhelm et al., 2021], it is not readily possible to port those changes to another engine to benefit from a proven method because two different engines may operate in totally different ways.

State-of-the-art peptide-centric search engines for peptide identifications can be classified in many different ways. One classification axis relevant to the difficulties in modifying engines for actual use with novel techniques is their license status (i.e. proprietary or not), and source status (i.e. closed vs. open, published vs. unpublished). Breakthrough deep learning algorithms have been proposed, but they cannot be added to proprietary search engines and released legally (except by such search engines’ authors themselves), slowing down research and making convincing experiments to demonstrated added value on top of some commonly used engines difficult. In addition, current open-source search engines do not have deep learning methods in mind and are designed in such a way that integrating machine learning models to their pipeline is difficult: for example, deep learning algorithms are designed for efficient batch processing, potentially at one specific stage of the process, whereas current engines are desgined to perform all processing steps across groups of inputs.

In this paper, we propose Pepid: a new, open-source search engine that is designed from the ground up with dry-lab research and deep learning methods in mind and, in general, to further research and development of tools and algorithms for peptide identification. Our search engine features a multi-phase, batched pipeline design. Each step in the pipeline is fully optional, uses configurable user function attachments, and supports a variety of user settings to customize computational schedules. We believe this is a more appropriate design for applications such as deep learning than the usual “end-to-end” approach of other options, at the cost of incurring some overhead that we show to be negligible overall.

We implement two common scoring algorithms and their extensions as well as a combination score function, and we show that Pepid is competitive in both runtime and identification rates. We believe that deep learning is already instrumental to achieving best-in-class performance in peptide identification, and that further research at the intersection of deep learning and proteomics is critical to next-generation peptide identification research. We provide experimental deep leanring methods with our source release to demonstrate how one might go about integrating such algorithms, although discussion of those details falls outside the scope of this paper.

## 2. System and methods

Existing search engines present various tradeoffs based on their implementation and design. We discuss a select few common examples and describe how Pepid addresses flaws in current choices. The selected engines showcase some orthogonal design characteristics that make them suboptimal for Pepid’s usecase (research into computational methods for peptide identification, as opposed to maximum identification rates or counts, and incidentally, throughput).

### 2.1 IdentiPy

IdentiPy Levitsky et al., 2018] is an open-source search engine written in python 2.7, using cython in parts to improve performance. Its main feature is its high configurability, providing users with custom extension points allowing staff with limited technical skills to provide and apply arbitrary python functions at limited parts of the pipeline. This paradigm also means the intermediary computation steps are exposed to the user, allowing more flexibility in how the pipeline operates and runs. It makes use of multiple processors via multi-processing and performs task dispatch on the unit spectrum level (that is, each query spectrum is individually processed by the next available process on a first-come-first-served basis with no task batching).

However, only a small set of extension points are supported, and the single task dispatch design means that operations with short computations and long preparation times (e.g. those leveraging GPUs for general-purpose computing, as would be desired for a deep learning-driven workflow, or those that should load data from the disk in chunks) limits what kind of functions can feasibly be implemented this way due to wallclock time concerns. Moreover, it is implemented in Python 2.7, which is incompatible with Python 3.0+ and is deprecated as of January 1st, 2020; conversion is not difficult but remains a burden on potential users. It also operates fully in-memory, which, combined with the multi-processing paradigm, makes it susceptible to the “memory explosion” problem - that is, as the task schedule is unpredictable and may use large amounts of memory, potentially exceeding available memory; this may cause the entire run to fail. It also means that the database size used in the search, as well as the query set used in any one run, are artificially restricted based on hardware availability.

### 2.2. X!Tandem

X ! Tandem Craig and Beavis, 2004] is an open-source version of the proprietary University of Washington’s SEQUEST engine Eng et al., 1994]. X Tandem is reasonably fast and efficacious, however it is designed to operate “end-to-end” for identifications, meaning there is no way to halt the process at arbitrary points. On the other hand, X !Tandem supports custom scores via a “score plugin” system. An example of a popular score plugin is the “k-score”, which is the same basic scoring method that was later implemented in Comet Tabb, 2015].

Score plugins in X !Tandem are developed in C++, which requires a certain technical sophistication from the programmer as well as providing a rather slow development cycle when working on iteratively improving the score function. By comparison, languages such as Python, Julia or Lua, which may feature slower operation at runtime, but provide better facilities for rapid application development, may be more appropriate for computational research purposes, since a better optimized version of the algorithm may replace the prototype after a novel approach is found to be advantageous.

X !Tandem also does not allow outputting more than the single best-scoring peptide match for each spectrum except when applying a filter on the score (e.g. there is no way to output an unfiltered best-10 score), which makes disambiguation with potentially close-in-quality matches impossible without further manipulation of the software, and score distribution analysis only possible with sufficient hardware to store all scores on disk, as well as temporarily in memory during X Tandem’s processing. This may also affect the performance of post-processing methods like Percolator Käll et al., 2007] as they may exploit the greater amount of available data if provided to them.

### 2.3 Comet

Comet Eng et al., 2012] is a reputedly fast search engine whose main scoring function is a function of the inner product between the theoretical spectrum of a candidate peptide and the empirical spectrum of the query input (where both spectra are represented as fixed-length vectors of mass to charge intervals to intensity values), normalized by the average of the same score across a window around each peak window. Comet also features a so-called E-value score, which is based on a regression over the score distribution’s log-histograms (i.e. the E-value score is an ordinary least-square approximation of the estimated log-CDF of the score distribution). Similar to X !Tandem, Comet is designed for “end-to-end” operations and does not innately support any modification to its runtime procedure. Unlike Xä !Tandem, it also does not support a plugin system for scores or other aspects of the computations.

Comet operates in a batch-oriented way, processing a user-specifiable amount of spectra at a time, thus bounding memory requirements, potentially at the cost of processing time. This design is more amenable to be modified to support heterogeneous compute capabilities. Indeed, Tempest Adamo and Gerber, 2016] is such a modified version of comet and can run on CPUs or GPUs using OpenCL for faster processing.

### 2.4. Others

A plethora of other search engines exist, some proprietary like PeaksDB Zhang et al., 2011], Andromeda Cox et al., 2011], or the Thermo Fisher version of SEQUEST Eng et al., 1994], and some open-source, such as Morpheus Kim and Pevzner, 2014a] or MS-GF+ Kim and Pevzner, 2014b], to give but a few arbitrary examples. To the best of our knowledge, beside potential additional issues related to licensing and availability of source code, all commonly used search engines share an “end-to-end” focus causing similar challenges to those described in the previous sections. Selection of the above search engines for longer exposition does not imply endorsement, and was not made on the basis of subjective of objective criteria other than the following:

- All three engines are open-source, allowing examination of internals;
- IdentiPy has inbuilt user extension points;
- Comet and X! Tandem are in common use in practice.

This provides a comparison background that enables us to estimate Pepid’s viability in practice (by proxy, using X! Tandem and Comet for comparison) as well as for fiexibility (especially by comparison with IdentiPy, which allows the most customization to workflow among engines we are aware of).

A qualitative summary of some search engines compared to the proposed Pepid engine is provided in Table 1.

**Table 1.** Qualitative summary of some search engines. Source is the status of the engine’s source code. Published Source means partial or snapshot source releases are available, for example accompanying a publication. Phased Runs means an engine can perform only a subset of its operations at a time and outputs artifacts that can be used to continue a run at a later time from where the engine left off. PSM Output qualifies if an engine can output only the top N outputs, or if it can filter based on score. *: SEQUEST is commonly used to refer both to the University of Washington and the Thermo Fisher versions of the software. The listing here is for the latter. *†*: The ldentiPy developers are in the process of updating their engine to Python 3 at the time of writing. *×*: Pepid outputs artifacts containing all scores that were above 0, and provides a condensed output with the Top N peptides for each spectrum by default. This is fully user-configurable.

| Engine | Language | Extensible | License | Source | Phased Runs | PSM Output | Speed |
| --- | --- | --- | --- | --- | --- | --- | --- |
| **X!Tandem** | C++ | Little | Permissive | Open | None | Filtered | Fast |
| **Comet** | C++ | No | Permissive | Open | None | Top N \| Filtered | Very Fast |
| **IdentiPy** | Python 2.7† | Somewhat | Permissive | Open | None | Top N | Slow |
| **SEQUEST\*** | C++ | No | Proprietary | Closed | None | Top N \| Filtered | Very Fast |
| **PeaksDB** | Java | No | Proprietary | Closed | None | Filtered | Slow |
| **Andromeda** | C# | No | Proprietary | Published | None | Filtered | Slow |
| **Pepid** | Python 3 | Fully | Permissive | Open | All | Any × | Fast |

### 2.5 Datasets

ProteomeTools is a dataset of synthetic human peptides. The dataset was generated using synthesized peptides based on the SwissProt database Bateman et al., 2020].

Due to corruption in the dataset (causing some of the archive files to fail to decompress), we only use the First Pool data (which is uncorrupted). We select only the higher-energy collision dissociation (HCD) data at a normalized collision energy (NCE) of 25 as this seems to produce the best quality results out of the available settings Zolg et al., 2017].

The One Hour Yeast Proteome is a dataset of non-synthetic yeast peptides with a baseline generated by the Thermo Fisher version of the SEQUEST search engine Hebert et al., 2014]. We use the “batched” dataset, which combines the result of all the hour-long runs.

The Massive-KB dataset contains spectra from real experiments and use a best-representative and consensus identification approach to provide a high-confidence peptide identity Wang et al., 2018]. We select the “human hcd in-vivo” subset and remove cross-digestion experiments (such as HeLa Trypsin-Lysc experiments) by excluding those entires whose provenance dataset contains keywords lysc, argc, or chymotryp. This dataset segment does not contain the ProteomeTools data which is included in the “full” Massive-KB dataset.

The ProteomeTools data provides an inordinately clean baseline using artificial peptides and pools designed to minimize mass confiicts, while the yeast dataset is a realistic real-world peptide dataset. Together, they allow us to qualitatively observe Pepid’s suitability in both settings, as each may provide a different yet useful setup for research purposes.

Properties for the datasets used in this work are listed in Table 2.

**Table 2.**
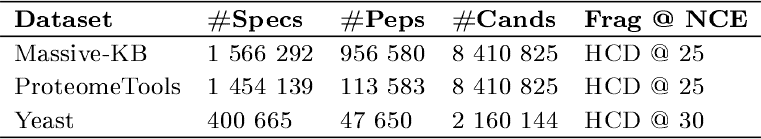
Features of the datasets used to evaluate Pepid. Peptides (#Peps) are unique pepide sequences, excluding modifications. Candidates (#Cands) are the count of Pepid-generated candidate peptides in total (i.e. including modifications, missed cleavages, and so forth) for the species database. Spectra (#Specs) are the total spectra in the query dataset.

**Table 3.** Spectrum generation average cosine similarity compared to Prosit. Performance on Massive-KB is on the held out set only for our method. Note that PredFull uses the pretrained model from predfull.com, which used a combination of proteometools, NlST [Stein, 2008] and Massive-KB data for training.

|  | Massive-KB | ProteomeTools | Yeast |
| --- | --- | --- | --- |
| PredFull | 0.629 | 0.573 | 0.320 |
| Ours | 0.632 | 0.616 | 0.320 |

### 2.6 Evaluation

Each search engine uses a different method for evaluation and reporting: for example, classical false discovery rate vs. q-value Käll et al., 2008], different formulae for their computation Elias and Gygi, 2007, 2009], etc. In order to ensure comparisons can be made, we use the same external evaluation method for each search engine rather than relying on their native tools. The code for evaluation is available in the script `gen_fdr_report.py` in the Pepid source release. We compute q-value using the FDR formula in Elias and Gygi 2007]. In addition, since FDR can be untrustworthy Gupta et al., 2011, He et al., 2015, Elias and Gygi, 2009, Matrix Science Ltd, 2010, Danilova et al., 2019, Jeong et al., 2012], we also report the false discovery proportion (FDP) using ProteomeTools’s groundtruth labels as computed using the same algorithm as for FDR.

### 2.7 Spectrum Generation

We present a spectrum generation model designed for integration into a search pipeline. Spectrum generation quality is benchmarked against Predfull, using code and model obtained on 2023-05-16, labelled as version 2022.05.19. The pretrained model was obtained from the PredFull Google Drive indicated in the PredFull github-hosted repository (https://github.com/lkytal/PredFull) Since this pretrained model was trained on ProteomeTools and thus may exhibit some overfitting compared to our model. The code was obtained from the master branch of the same github repository. We choose to compare to PredFull rather than alternatives (like Prosit or DeepMass) for the following reasons: first, PredFull was previously compared to other models and found to perform better even with only the theoretical peak sequence. Second, our method also performs full spectrum prediction, making the comparison more directly appropriate. Third, PredFull is easily reproducible and is still maintained, whereas other models like Prosit have not seen an update since 2018 (as of the time of writing) and can no longer be loaded with the currently available versions of existing dependencies.

## 3. Algorithm

Pepid is designed to operate in restartable, customizable phases. Each phase can be customized both by user parameters describing behavior settings (for example, how to multiprocess the program), or by custom (i.e. fully user-written), user-specified functions (the score function, the peptide candidate generation function and the rescoring function, for example, are all user-provided, with sensible defaults implemented by Pepid). Pepid uses python’s dynamic capabilities to read the name of the functions provided in a configuration file and dispatches work through these functions as appropriate during operations. In addition, users may elect to perform a subset of phases before directly examining or modifying the data, and then proceeding with remaining phases.

### 3.1. Inputs and Outputs

User configuration to the pipeline is composed of a single file: a user configuration document in an INI-like format compatible with the python `configparser` module, for which a default configuration set is provided in the `data/default.cfg` file in the release distribution. The default file features explanatory comments for each configuration option to guide modifications. The set of query spectra as specified in the configuration file must be in the Mascot Generic Format (mgf) and the database of candidate proteins must be in the protein fasta format. Pepid’s output is a database of processed queries, candidate sequences (generated in-silico from the fasta input database), and peptide-spectrum matches (PSMs) as SQLite databases. An optional pipeline step also extracts the top *N* (for some user-selected *N*) matches per query spectrum and outputs them as a tab-separated value (tsv) file. Afterward, the pipeline optionally runs percolator Käll et al., 2007] or a custom random-forest-based rescoring method on the output, and can generate a graphical report about identification performance using the target-decoy approach, false discovery rate (TDA-FDR) methodology, both after and before rescoring.

### 3.2 Pipeline

The search engine features a pipeline design composed of 6 steps: Query and Database Processing, Input Postprocessing, Search, Report, Rescoring, Final Report, which we explain in more details below. The pipeline is graphically illustrated in Figure 1A.

**Fig. 1:**
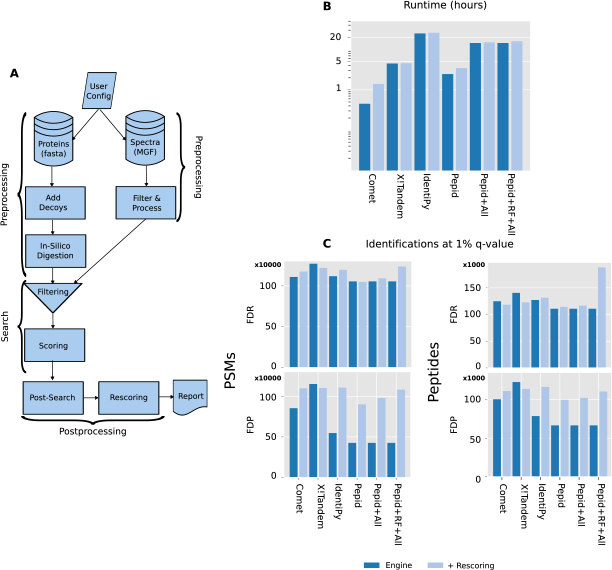
Comparison between Pepid and commonly-used search engines on ProteomeTools first pool (1 458 831 spectra). RF: using the random forest-based rescorer instead of Percolator. All: using both the spectrum generator and peptide length prediction models. IdentiPy cannot process the spectra witin resource limits, so the file had to be split 10-ways, and the process run once per split. X Tandem is only capable of outputting a single top PSM when not applying a score filter, other engines output the top 10. A: High-level processing pipeline steps of Pepid. B: Runtime performance. C: Count of identifications (for FDP, as per ProteomeTools baseline) at a q-value of 1% (PSMs on left, peptides on right, TDA-FDR on top, FDP on bottom).

In the Query Processing step, the input query file is processed in a more suitable format for further operations and is stored in a SQLite database. In the Database Processing step, peptides are generated from a fasta protein database. The resulting peptides are saved in a SQLite database. User-specified functions are used to process the queries as well as to generate peptide candidates.

In the Input Postprocessing step, user-defined functions are applied to the processed queries and peptides. The user is thus able to insert arbitrary data into the database.

Besides the arbitrary post-processing function, the user-specified spectrum prediction function is applied to the database candidates. The default spectrum prediction function simply generates the theoretical b and y ion series for the candidate modified sequence for each charge level indicated by the user. The output spectrum object can be an arbitrary python object and will be inserted as a binary blob into the database.

In the Search step, a user-specified scoring function is applied to a set of queries and a set of candidates for each query. The candidates are a subset of the entire database that matches a user-specified tolerance threshold around the mass of the query precursor neutral mass, as is usually done in other engines to restrict the set of candidates to search against.

The Report step prepares a graphical report containing a mix of performance metrics based on the TDA-FDR methodology, along with score distribution data plotted against basic PSM statistics, to help identify potential biases or weaknesses in the score function. The Report step also outputs a serialized artifact containing the statistics computed during the report generation for further analysis. This artifact can also be used to generate comparison plots using the `pepid_compare.py` script which is discussed further below.

The Rescoring step applies an arbitrary rescoring function, which is Percolator by default, to the results.

The Final Report step produces the same report artifacts as the Report step, only after the rescoring function is applied.

Pepid phases are reusable and optional throughout its process: unlike other search engines, the intermediary artifacts remain between each step and the software can rerun only the steps of interest to the user. This also enables the user to perform part of the steps, then use custom software to modify the resulting artifact, then proceed with additional steps, without Pepid having to know anything about the intermediary user processing scripts, granting additional pipeline fiexibility beyond the basic pipeline proposed by the Pepid system. It also allows users to perform perhaps lengthy processing (especially for peptide generation -that is, in-silico digestion by more complex algorithms and or spectrum for candidate peptides) only once and to reuse this database for as many query searches as required. Another potential use case is to first generate the preprocessed database on one machine (or multiple machines and then performing a merge step), and making that database available on multiple hosts for search across a cluster (before performing a synchronized final merge of these databases). This design also affords some robustness regarding computational disruptions.

Pepid also provides additional utility scripts: `pepid_compare.py` takes two report artifacts and generates a combined report visualization that makes comparing search results between two conditions convenient. `pepid_files.py` outputs the paths to the files generated by Pepid as part of the search process, given the class of files of interest (for example, just report artifacts or just database artifacts).

### 3.3. Preprocessing

The query preprocessing step simply adds useful metadata, such as the count of peaks in the spectrum and the precursor neutral mass, to the reformated input data, and places it in a database for more efficient further processing. The user can specify criteria used to filter query spectra for quality, such as the peak count range that is allowable and the maximum and minimum precursor mass.

On the other hand, the database processing step is more involved. Its main substeps are as follow: In-silico Digestion, Filtering, Post-translational Modification (PTM) generation, Deduplication.

In the first step, a user-specified regular expression (regex) is applied to the protein sequences in the database to extract “digested” peptides. The regex specified by default corresponds to tryptic digestion using the EXPASY rule. Users can additionally specify maximum missed cleavages. The resulting peptides are then filtered by user-specified minimum and maximum mass and length for the sequences. Next, user-specified maximum variable modifications, variable modification types, and fixed modifications are applied against the peptides kept from the previous step, triggering a refiltering for mass. Finally, duplicate peptide-modification tuples are merged by joining the source protein identifiers to retain a single entry for each unique such tuple.

### 3.4. Search

The search procedure proper first selects a batch of candidates matching a certain tolerance window as specified by the user around the precursor neutral mass of each individual query spectrum, then applies a user-specified scoring function to the batch of candidates for the input query spectrum. Any resulting match scoring more than 0 is kept in the final results database (it is the responsibility of the score implementation to adapt to this 0 cutoff based on other user settings to achieve the desired behavior). The complete artifact takes the form of the query spectrum identifier (the title provided in the input mgf file), the source protein identifier (the protein identification line from the fasta files, or multiple lines merged with semicolons if duplicate peptides were generated in the preprocessing step), the score, and an arbitrary python metadata artifact as a binary blob. It is up to the scoring function implementer to ensure that the artifact contains all the data necessary to apply the rescoring function if such a step is desired later.

The default scoring function provided by Pepid is a combination of an implementation of the Xcorr function based on the Comet codebase and an implementation of the Hyperscore function based on the X! Tandem codebase, a strategy that has been shown in the past to improve search result qualities [Shteynberg et al., 2013]. We also modify our Xcorr and Hyperscore implementations in two major ways: first, we expose many implementation details of those algorithms as user modifiable parameters in our configuration file, and second, we expand in multiple directions upon those algorithms. Our extensions are as follows:

- We generalized comet’s so-called “fianking peaks” feature to support using arbitrary amount of intensity bins on either side of the current bin during spectrum correction;
- We implement a gaussian kernel and a generalized version of the original comet kernel using the exponential falloff 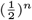 to weigh intensity values *n* bins away;
- We implement dynamic bin boundary settings based on ppm calculations at the level of individual spectra, supporting 3 operating modes: in “max” mode, the highest m z in the spectrum is used to compute a m z difference equivalent and is the basis for setting the distance between bins; in “precursor” mode, calculation proceeds as in “max” mode, but operates off of the percursor neutral mass; in “bins” mode, bin boundaries are iteratively computed from the maximum allowable mass down to the user-provided minimum mass interval using the distance metric computed as in “max” mode.

We similarly apply modifications to the hyperscore algorithm as follows:

- We keep the top N best-matching sequences, with user-provided N and user-provided series selection criterion (i.e. answering the question “best N what?”): “charge” instructs Pepid to keep the top N charges, counting matches across all series for the same charge level; “series” does the converse (selecting the two best series across all charge levels); “both” selects the best matching vectors (i.e. series-charge sequences).
- We make the theoretical spectrum weighing scheme used by X ! Tandem (corresponding to the input parameter ‘refine, spectrum synthesis’) user-settable and optional.
- We add a score reformulation option that modifies the model to assume that the sum of intensities (all sequences) is caused by the selected best-matching sequence(s) alone (rather than the joint of all selected sequences as in the original X Tandem model).

We show that some of those modifications can greatly improve peptide identification rates in Section 4. Additionally, we deliberately abstain from implementing the X! Tandem hyperscore protein-level peptide score adjustment and the Comet E-Value scoring function because it is well known that both methods defeat appropriate TDA-FDR evaluation (for example, it was noted by Zhang et al. 2012], Gupta et al. 2011], Matrix Science Ltd 2010], Jeong et al. 2012] and indirectly observed in Elias and Gygi 2009]).

The uncombined version of each scoring function (i.e. Xcorr and Hyperscore) are also made available to users by default by selecting the right function in the configuration file.

### 3.5. Input Postprocessing

In Input Postprocessing, a user-specified function receives an input python dictionary-like object describing the corresponding entries in the database that are to be user-processed, and outputs an arbitrary python datum that is then serialized as binary and saved in the database in an extra column. It is up to the user to implement the correct processing functions at the correct stages of the pipeline to make use of this data, or to rely on the primitives already provided by the engine.

In previous work, we added a deep-learning length prediction system as an input postprocessing step applied to the set of queries. The length prediction module adds a vector of probabilities corresponding to the predicted probability of the peptide being of a given length as a metadata item to each query spectrum.

We also add a deep learning-driven spectrum prediction input postprocessing module to the input postprocessing step for candidates. For each candidate, the deep learning module predicts a full spectrum (i.e. both the m/z and the intensity across the entire spectrum, not just the intensity for preselected masses as in Prosit Gessulat et al., 2019] or DeepMass Tiwary et al., 2019]), which is added as a metadata item. We do not use this module as the main spectrum generation function because, as with Gessulat et al. 2019], Wilhelm et al. 2021], we find that our predicted spectrum is better used as a feature for a rescoring algorithm like Percolator.

### 3.6. Postsearch

A postprocessing phase following the search, dubbed “postsearch” to avoid ambiguity with other postprocessing phases, is applied before the rescoring phase. When using the length prediction module mentioned above, this phase can add various computed features based on the PSM characteristics to the metadata for each search result. This design was chosen mostly as a demonstration of the pipeline’s fiexibility, as it could be more efficient to only perform this step as an out-of-pipeline operation as we do to append groundtruth labels to the data (described in implementation), so as to only generate predictions for those entries that will be used for rescoring. Nevertheless, this design allows examining the results to drive potential insights and further peptide identification research.

### 3.7. Rescoring

The rescoring module runs the user-specified function on the input results rows. The default rescoring function generages a Percolator INput file (PIN), and uses Percolator for rescoring, converting the Percolator OUTput file (POUT) at the end of the Percolator rescoring process into a Pepid output tsv. Alternatively, Pepid also provides a random forest-based rescoring method.

We also provide an experimental random forest-based rescorer that is pretrained on Massive-KB Wang et al., 2018] (a dataset of non-synthetic human peptides with a consensus-based ground truth), in the style of the original formulation of PeptideProphet [Ma et al., 2012]. We show that this rescoring method appears to generalize as it successfully rescores both ProteomeTools and One Hour Yeast Proteome search results beyond what Percolator is capable of at both the FDR and FDP level despite no finetuning being applied.

### 3.8. Reports

The Report and Final Report steps are the same, except at different stages of the pipeline: the Final Report operating after rescoring. The result of this step are text file artifacts containing data about PSM quality across FDR thresholds and graphical artifacts summarizing the results (see Figure 4 for an example of the output graphical artifacts).

### 3.9. Out-of-pipe ine Operations

The phased, pipeline design of Pepid allows interrupting the process at any time and using arbitrary functions to alter the contents of the databases used to store the queries, candidates or search results. We showcase this capability by implementing such an out-of-pipeline insertion of fields from the mgf file that aren’t otherwise recorded by the query processing step (namely the ground truth sequence from ProteomeTools used exclusively for FDP evaluation, which is introduced in the mgf data by the SEQ= field).

## 4. Implementation

Pepid is implemented in Python 3. It is accelerated using a custom multiprocessing module available under pepid_mp.py in the source repository, which allows greater fiexibility for reporting and debugging than usual python options such as multiprocessing. The scoring functions are also accelerated using numba, and extensive use of numpy and vectorization provides additional speed advantages. For the experimental deep learning facilities, pytorch is used for both training and inference.

The sqlite database is used as the backing store for the artifacts at each stages. During search, heavy sharding is used to ensure maximum throughput. This means that Pepid is highly sensitive to the storage medium and properties for the destination of the artifacts. To avoid performance degradation due to operations on a large SQLite database, each process creates a new shard whenever a user-specified result count threshold is exceeded.

The condensed output generation (i.e. the output to a human-readable tsv file) and the PIN generation steps can be very slow if operated sequentially, so they are multiprocessed as well. To avoid data races, a file lock is used to negotiate write access to the appropriate file. This is sufficient to speed the process up to 2000% compared to sequential operations.

In the current version, Pepid relies on unix-specific facilities to perform both the multiprocessing (UNIX sockets are used for communication), as well as for the file lock facility.

### 4.1. Search Parameters

Search parameters for the cross-engine comparison using ProteomeTools were selected to match as well as possible between the search engines used, while enabling any special method available to each engine as would normally be enabled in a real search. A summary in the Pepid configuration format is provided with the software. For the yeast search, parameters for Pepid were optimized so as to represent realistic performance statistics for more accurate comparison on the basis of wallclock time.

While Pepid supports rescoring by Percolator or by a partially pretrained random forest method, we only compare engines with percolator due to the pervasiveness of this method across established pipelines. We show the results of the pretrained random forest method compared to previously shown Pepid results to demonstrate generalization and increased identification rates separately. Due to its pretrained nature, this random forest approach cannot equitably be applied to other search engines.

### 4.2. Length Feature Extraction

When the length prediction module is in use, several features are computed in postsearch and added to every PSM from the search step. In particular, we compute the difference between the most-probable predicted length and the candidate length, the predicted probability that the peptide present in the spectrum is of the candidate’s length, the difference between the probability of the candidate’s length and the best, worst, next best and next worse length probability. Those features are simply extracted during the rescoring step and used as extra features in the Percolator-based pipeline, for example.

### 4.3. Spectrum Generation

We developed a deep learning-based approach to full-spectrum prediction. To the best of our knowledge, the only previous work to have ever successfully used this approach was PredFull Liu et al., 2020]. Our approach differs in several aspects: first, we use fully standard convolutional neural network instead of PredFull’s mix of “Spike and Excitation” units and multi-scale convolutions. Second, we output a predicted spectrum for each charge level (equivalent to using 5 different networks that heavily share weights). Third, we constrain the output spectrum so that any intensity less than 1 · 10^*−*3^ is dropped to 0 so as to obtain a sparse spectrum (we find that resulting spectra are usually very sparse, at around 1-3 non-null peaks per 1000 bins, with a total of 50000 bins, i.e. an average of up to 150 peaks per generated spectrum). We do this mostly due to storage constraints when applying the model during peptide search, although this also acts like a noise removal pass.

The model’s architecture and input processing are presented in Figure 2. It is can be understood as an encoder (a convolutional layer processing the whole sequence at once), 5 processing stages (using convolutional residual layers with a kernel size of 1), and 5 decoders (one per charge level), each independently transmuting a size 1024 latent representation into a size 50000 predicted spectrum corresponding to each input in the sequence, which is then averaged over those sequence elements. The result is a predicted total spectrum in rasterized (i.e. a vector for which each bin has a 0.1 Da width) format for each charge level. This design allows the model to be used efficiently to preprocess a database of candidate peptides, without having to repeat this relatively slow prediction process when iterating upon scoring mechanisms or parameters.

**Fig. 2:**
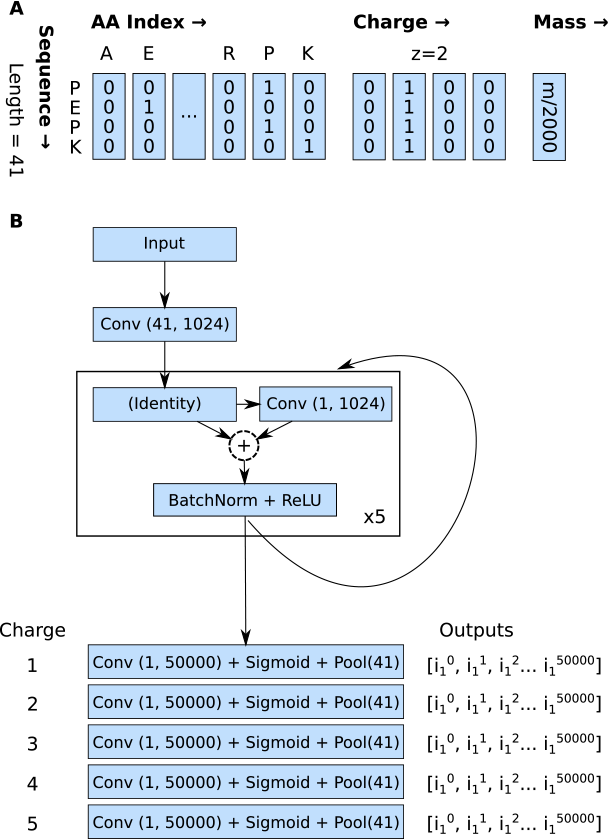
Architecture and input transformation for the Spectrum Generation model. A: The input peptide sequence is encoded similarly to PredFull Liu et al., 2020]: one-hot peptide sequence, one-hot charge, and mass divided by a fixed scaling factor (2000 here). B: The model architecture is composed of straightforward convolutional layers (kernel size, embedding output size). The final pooling layer averages over the sigmoid outputs across each of the 41 input sequence elements, in effect gathering the consensus from the spectrum implied by each amino acid. The output is 50000 intensity levels for each output charge level. The dashed circle with + represents elementwise addition.

We train the model using the Massive-KB data to avoid potential overfitting vs ProteomeTools as much as possible, and we show some results on ProteomeTools and the Yeast dataset to demonstrate that the resulting model is generalizable. Finally, we apply it during candidate post-processing and store it using the compressed sparse row representation using the SciPy module, version 1.10.0 Virtanen, 2020].

### 4.4. Rescoring

Beside the classical Percolator-based pipeline, we also present a ground truth-based, pretrained, random forest rescorer using the Massive-KB dataset so as to avoid overfitting as much as practical. To demonstrate its capabilities, we measure performance both for ProteomeTools search results with Pepid, and Yeast search results, both on the basis of FDP and FDR identifications over common value ranges. The model is implemented in scikit-learn Pedregosa et al., 2011] version 1.3.3 as a pipeline composed of a standard scaler followed by the random forest. The model is trained in a 10-way cross validation setting and the best-performing model on the basis of separation between ground truth and other hits is saved (along with the scaler). The model and scaler are then loaded and applied to the search results to rescore (i.e. those of ProteomeTools and Yeast in this case), and a second random forest-based rescorer is trained to identify high-scoring vs low-scoring targets (where low-scoring means those with a score below the median score for the collection of PSMs including decoys). This is used as a fine-tuning phase, and the output probability is averaged with the pretrained ground-truth-based random forest probability outputs to generate the final score. The fine-tuning is required to adapt the score to some different experimental conditions, such as search parameters or species under investigation, as we show using the Yeast dataset.

### 4.5. MGF Field Insertion

To compute FDP-based metrics, a separate module (`pepid_mgf_meta.py`) is provided in the Pepid distribution. It takes the configuration file and a field name and inserts the field in each entry of the mgf file into the corresponding entry in the queries database (if the mgf entry was present in the database). This serves also as a demonstrate of working with out-of-pipeline processes to achieve even higher fiexibility in overall processing.

### 4.6. Results

Pepid was compared against other search engines to demonstrate that it forms a solid baseline, without which its utility as a research platform would prove limited. It was also compared against itself using different datasets to show how Pepid scales based on dataset size in terms of runtime.

For the comparison between search engines, runtime and identification rates are considered. We use the ProteomeTools dataset as a comparison basis because it contains a large set of clean spectra which, combined with our hardware capabilities, serve to better explore real-world performance metrics than would a smaller dataset.

### 4.7. Performance

Performance results for the spectrum generation task on the whole spectrum are presented in Figure 3. We note that our model is trained only on the Massive-KB data, while the PredFull model is trained on Massive-KB, ProteomeTools, and Nist peptide datasets. Despite this, our model performs slightly better on Massive-KB, and much better on ProteomeTools. The indistinguishable, yet low, performance on the One Hour Yeast Proteome dataset is likely due to reaching a ceiling due to the dataset not having a real ground truth (it uses Sequest search results instead) and the spectra in the dataset not being quality-selected, such as by originating from synthetic peptides, or being collected from consensus as in ProteomeTools and Massive-KB respectively. Another potential discrepancy is the difference in NCE. Higher NCE typically results in more relatively pronounced low-mass peaks. Since our model is trained on Massive-KB, it could be underestimating intensities at lower m z values compared to the expected yeast spectrum.

Runtime performance was evaluated on a machine with the following relevant hardware:

- RAM: 192GB
- CPU: Intel(R) Xeon(R) Gold 6130 CPU @ 2.10GHz (64 pseudo-cores)
- GPU: 1x Nvidia GeForce RTX 2080 (12GB VRam)
- Disk: Intel Corporation NVMe Datacenter SSD 3DNAND, Beta Rock Controller] Model: SSDPE2KE016T8
- OS: CentOS, kernel version 3.10.0

**Fig. 3:**
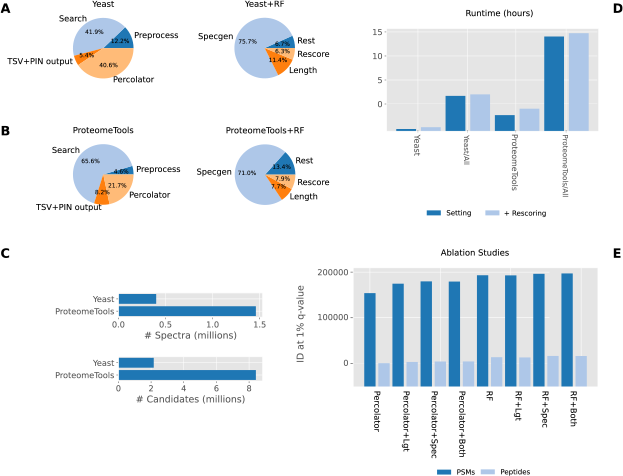
Performance scaling data for Pepid. Lgt: length predictions. Spec: deep learning spectrum generation. Both: Lgt+Spec. RF: random forest-based rescoring. A,B: Runtime breakdown by phase per dataset. Right compares the runtime for computing all machine learning features vs non-ML (“rest”) tasks. C: Indicative database sizes (queries and candidates). D: Overall runtime comparison. E: PSM and peptide identifications depending on which of Pepid’s modules are in use.

**Fig. 4:**
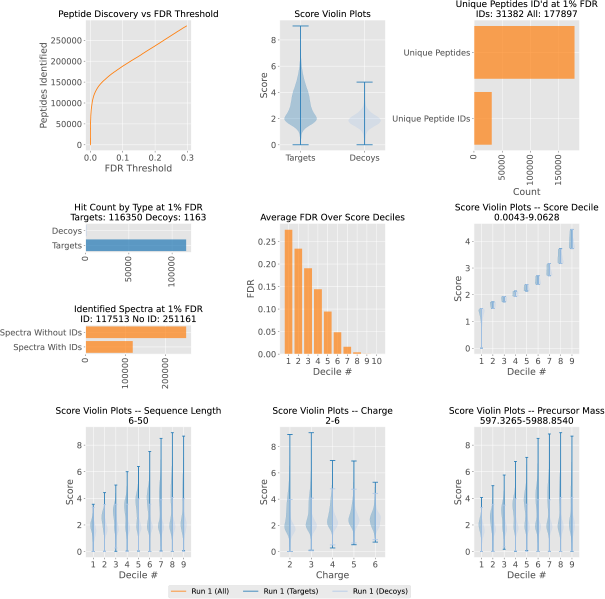
Example unretouched analysis output from the default Pepid “report” module (run on the One Hour Yeast Proteome data).

Figure 1B summarizes the runtime performance of various engines on ProteomeTools so as to demonstrate Pepid viability in a realistic workload, while Figure 3 compares Pepid’s scaling performance on our datasets and demonstrates that the added fiexibility in the hyperscore and xcorr implementations can be beneficial for identifications in some contexts. The combined xcorr and hyperscore scoring function is used in all experiments. We match our parameters to that of other engines as closely as possible for the cross-engine comparisons. For the Pepid performance scaling experiments, we vary the parameters to show the relative performance of our engine in these scenarios when performing a realistic search function. We find that for both dataset, although especially the yeast dataset, we can improve upon results provided by plain xcorr or hyperscore by using our algorithm extensions, as shown in Figure 3E. We make use of Pepid’s phased design to quickly iterate upon search settings or parameters to find the best choices without having to incur input processing and input postprocessing costs (or even search costs when iterating on just pre-rescoring features), while we had to optimize the results the “hard way” for the other search engines, resulting in a slower and more painful iteration process. Due to the relatively painless nature of Pepid development, exposing more parameters and function variants of the hyperscore and xcorr scoring functions was straightforward.

Figure 3A,B demonstrates that while the phased design of Pepid incurs an overhead for preprocessing and output, this overhead diminishes quickly when dataset size increases, allowing search and rescoring wallclock time to dominate the overall process duration. This also appears to be true for the overhead of the machine learning algorithms, although they remain quite expensive to run overall. Thankfully, the phased design employed by Pepid means they only need to be run one per query set or candidate database respectively, and the resulting computed features are “portable” (i.e. they can be reused in different settings without regeneration).

As presented in Figure 1A, despite Pepid keeping all results in the final database on disk, while other engines typically work in-memory and maintain a limited set of top PSMs, it remains competitive in runtime compared to other common search engines. We show that Pepid is faster than currently used search engines, demonstrating that the design’s impact on overall speed is minimal in the current iteration. On the other hand, space usage may be an issue, as storing all the data for the One Hour Yeast Proteome search results takes about 160GB on disk, while for Proteometools, 2.8TB are required. However, deep learning-based methods increase runtime considerably at this time. Though that may be the case, the identification performance impact of our methods are significant in the settings tested in this paper, despite our approaches leaving space for refinement across the board (both for runtime performance and for metric performance).

In Figure 1C, performance between the selected search engines is compared, showing that Pepid performs well compared to other engines in similar parameter settings. A baseline of this level is important as a starting point for further research, as it shows that Pepid works reasonably well without a significant algorithmic update. We believe that this demonstrates once again that Pepid is a suitable tool for further research in peptide identification.

### 4.8. Visualization

It is well known that the hyperscore and xcorr scores are biased, notably for peptide length (i.e. they tend to generate higher scores for longer peptides) Wang et al., 2015, Hubler et al., 2019, Jeong et al., 2012] or charge Granholm et al., 2012]. For peptide identification algorithm research, it is important to have at least two basic metrics: accuracy (that which we aim to optimize, e.g. peptide identification rate) and bias (which we would like to minimize). Pepid currently offers basic visualization aids to expose common sources of biases relating to precursor charge, precursor mass, peptide length, identifications at a selected q-value FDR threshold at the peptide level, spectrum level, and PSM level, and identifiations over FDR, among others. These tools can be used to quickly identify biases, distribution skews, algorithm performance, etc. In Figure 4, a report plot for the xcorr function on the One Hour Yeast Proteome is shown. As previously reported in the literature, we find peptide length bias in the score distribution, demonstrating the potential usefulness of this tool for further development.

## 5. Discussion

We have developed a research platform for peptide identification research that is competitive with state-of-the-art methods in runtime and identification rate performance. Overall, Pepid showcases a design that is specifically oriented toward bioinformatics research and is suitable, in the authors’ experience, to combination with deep learning methods, all the while retaining runtime and identification performance in line or superior to commonly used “search-only” engines (that is, engines designed for throughput, not for modification and research).

We have shown that Pepid can easily be adapted to include deep learning methods in the search process, demonstrating the added value of the phased design. The custom function extension points displayed their value both during the development process (allowing simple configuration modifications to easily compare performance in different conditions) and the research process (allowing the easy extension of the Pepid run to use deep learning models). Furthermore, we have shown that the inclusion of deep learning tools for parts of the peptide identification pipeline can greatly increase identification rates across the board, and that those deep learning tools can be generalized to different species and, to some extent, experimental settings.

Currently, Pepid does not feature a score filtering system. That is because in this initial design, Pepid was developed for peptide identification research first, where we would like to keep as many search results as possible so that we may identify where search algorithms show weaknesses. This data is also usable to train deep learning models in various ways, where the presence of low-quality search results is important to help models explore a richer and more accurate data distribution. In a future update, we plan to address the disadvantage this brings in terms of space constraints.

Pepid’s phased design allows it to naturally operate in a distributed environment even though no such feature is officially supported at present; a simple scheme to achieve this is as follows:

- Perform input processing on a single node;
- Migrate the database artifacts to each computer that will be used for search;
- Perform only the search step on each computer, providing a mgf file subset appropriate for each;
- Migrate the resulting search databases to a single node;
- Perform output, report and rescoring on that node, as desired.

In a future update, more attention will be given to user-friendliness, for example by providing easy name association for enzymes instead of requiring the user enter a regular expression directly. In similar veins, An official utility script akin to the comparison and filepath generation scripts described previously will be provided to make other operations, like distributed processing, more convenient. In addition, although Pepid’s preprocessing function can be changed by a user’s arbitrary function instead, better defaults will be provided, for example expanding modification support and fragmentation types.

Finally, Pepid currently relies on UNIX facilities. While it may or may not operate under Windows using WSL2, a future release may get rid of the UNIX-specific facilitie in favor of a cross-platform option.

Regarding the newly introduced models, the spectrum generator is not aware of NCE during generation and therefore may require postprocessing-based adjustment, or the integration of larger datasets with a range of NCEs, to work better in different NCE scenarios. The lack of quality synthetic or consensus data of sufficient size, especially for non-human peptides and for non-tryptic digests, makes accurate evaluation of this, and the length prediction model, difficult. Similarly, the lack of reliable ground truth peptide identities makes proper FDP evaluation (to qualify FDR estimation quality) less than ideal, a problem that also affects other search engines as shown in our figures (for example, Comet appears to find fewer peptides than X ! Tandem with 1% FDR control, but far more at the 1% FDP level. This shows that X Tandem’s algorithm causes underestimation of FDP by the TDA-FDR method, and that Comet should be preferred in this example. The question to solve is: how can we obtain such confidence when we have no access to ground truth peptides?)

Our search engine is publicly available online at https://github.com/lemieux-lab/pepid.

## 6. Acknowledgement

Funding for this project was provided through the Fundamental Research Grant program from the Institut de Valorisation des Données (IVADO)^1^ .

## Footnotes

1 There is no number or identification associated with this grant

## Notes

### Competing Interest Statement

The authors have declared no competing interest.

